# Modelling eNvironment for Isoforms (MoNvIso): A general platform to predict structural determinants of protein isoforms in genetic diseases

**DOI:** 10.1101/2022.04.21.489045

**Authors:** Francesco Oliva, Francesco Musiani, Alejandro Giorgetti, Silvia De Rubeis, Oksana Sorokina, J. Douglas Armstrong, Paolo Carloni, Paolo Ruggerone

## Abstract

The seamless integration of human disease-related mutation data into protein structures is an essential component of any attempt to correctly assess the impact of the mutation. The key step preliminary to any structural modelling is the identification of the correct isoform onto which mutations should be mapped because there are several functionally different protein isoforms from the same gene. To handle large sets of data coming from omics techniques, this challenging task should be automatized. Here we present our code MoNvIso (<u>Mo</u>delling e<u>Nv</u>ironment for <u>Iso</u>forms), which pinpoints the correct isoform associated with the mutation of interest and builds a structural model of both the wild type isoform and the related variants starting from the name of the gene and the list of mutations of interest.

## Introduction

The spatial and functional diversity of the 20,465 protein-coding genes^1^ (https://www.ensembl.org/) in humans is dramatically augmented through alternative splicing that result in an enormous number of potential protein isoforms. Exact numbers are not fully known but common estimates for total isoforms are in the 10X range (245,000 transcripts in https://www.ensembl.org/). Alternative splicing can result in isoforms with relatively subtle changes to those that vary enormously in their structure, function and subcellular spatial expression^2^.

Indeed, most functional (and dysfunctional) biochemical processes are impacted by the expressed isoforms, which feature distinct functional roles. Examples of this complexity are the neuroligin and neurexin families, which perform synaptic regulatory functions that are surprisingly isoform specific^3,4^. This complexity is compounded by the addition of genetic variants which can directly influence the protein sequence of any affected isoform. Genetic variations can also affect the splice mechanisms and change the isoforms directly^2^, but this is not addressed in this study.

Further information, key to our understanding of a genetic disease, is the availability of three-dimensional structures of a protein. The structure of many human proteins is now available by accurate (yet time-consuming)^5,6^ experimental techniques (such as X-ray diffraction, NMR and electron microscopy^7^), complemented by fast (and more approximate) computational predictions^8^, from homology modelling^8^ to deep learning techniques such as AlphaFold^9^, based on the three-dimensional structure of evolutionarily related template protein(s)^8^. However, unfortunately, all of these methods do not provide the isoforms most likely involved in the process of interest.

Here we present a computational platform able to select specifically the best isoform in the specific case of genetic diseases. The presence of a variable number of protein isoforms makes it challenging to assign each mutation to a specific position in the protein sequence, which frequently hampers a reliable assessment of the impact of the genetic variations (including disease relevant mutations^10,11^) on the correct isoform. In other cases, a mutation is observed that is relevant to a specific isoform, but the databases reporting mutations related to a particular genetic disease usually lack a reference to the specific isoform.

Our pipeline systematically identifies the isoform(s) relevant to a gene level mutation or variant on which the pathogenic mutations retrieved from the literature, or from the clinic, can be correctly and efficiently assigned. The key step of the determination of the appropriate isoform is achieved by combining the information on the mutations with the coverage of the sequence by available templates.

### The MoNvIso (<u>Mo</u>delling e<u>Nv</u>ironment for <u>Iso</u>forms) pipeline

The general workflow of MoNvIso, described stepwise below is also summarised in Figure 1. The input is composed of a list of human genes and by a list of mutations in each gene.

**Figure 1.**
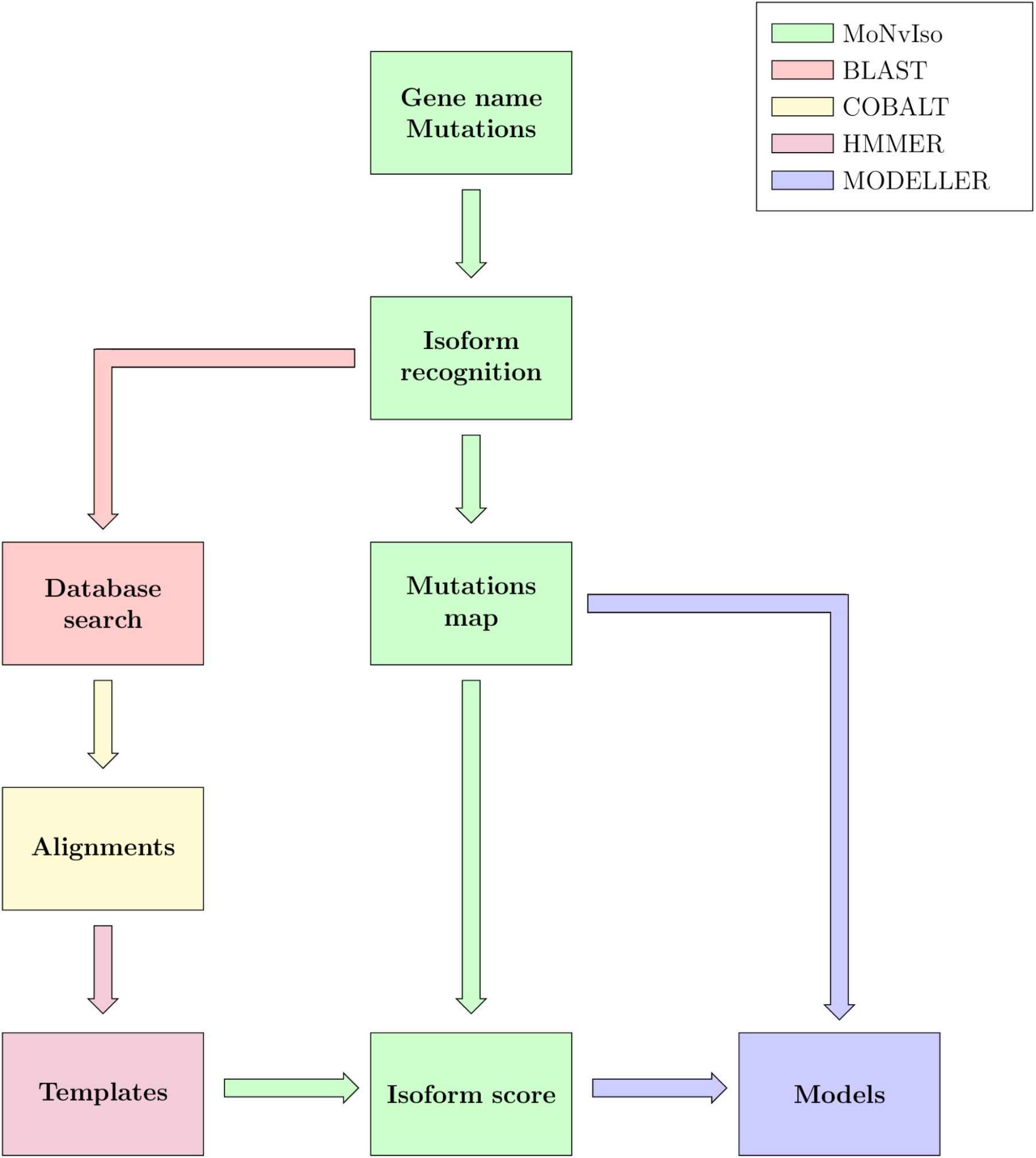
MoNvIso’s flowchart.

#### Step 1

As a first step, MoNvIso checks the provided list of gene names and single point mutations. The mutations can be inserted in the input file according to different formats: three-letters or single letter names for the amino acids. Additionally, spaces and tabs are also accepted to simplify the creation of the list by the user. Every gene name is searched against the Uniprot^12^ database, the results are stored in two files, namely *uniprot_sprot.fasta*, which contains the canonical isoforms according to the classification of Uniprot, and *uniprot_sprot_varsplic.fasta* collecting the remaining listed isoforms.

#### Step 2

Each isoform is processed according to a standard procedure: After a search for homologous sequences performed using the BLAST^13^ API, the generation of the associated multi sequence alignment (MSA) is generated using COBALT^14^, a HMM (Hidden Markov Model)^15^ is generated by HMMER version 3.3.2 (http://hmmer.org/) and this is used to find relevant templates in the PDB database. The 10 most similar sequences for the identified PDB structures are downloaded along with the chains necessary for the homology modelling which are extracted as separate files. The extracted structures are cleaned from water molecules, ligands, disordered atoms and non standard residues, then aligned to the MSA. The resulting structures are ranked by resolution and sequence identity to find the most appropriate templates that exclude crystals with poor resolution or with sequences that are very different from the original query (see Section Limitations). A further criterion is applied by calculating the coverage of the input sequence by the sequences of the templates. This is done to select the minimum number of templates necessary to model the highest percentage of the target sequence, avoiding superpositions, as much as possible. The total selection function is a sum of two terms:

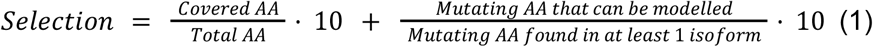

The first term of the selection function accounts for the availability of crystallographic data defined as the number of amino acids (AAs) that are covered by a template over the total number of AAs constituting the isoform

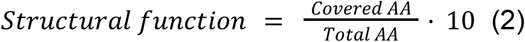

The list of mutations enters in the second term, which pinpoints the best isoform to be modelled. Our program maps all mutations onto the appropriate isoform and increases by one the numerator, ***Mutating AA that can be modelled***, if the mutated residue can be correctly located in the isoform sequence. The contribution of matched mutations to the selection function is evaluated as follows:

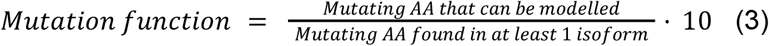

For each gene and each isoform, the resulting ***Selections*** are reported in a file called *report.log* collecting information, such as mutations not mapped, mapped mutations and on which isoform, the selection functions, and the isoform most suitable to be modelled.

#### Step 3

Structural models for the selected isoform are then created based on the sequence alignment obtained in the previous step by using the MODELLER program^16^. Regions not covered by the templates are not considered. The best models are then selected using the DOPE score^17^. In our case we select the best single model. The modelling of the variants is then performed by taking the MODELLER input file containing the WT sequences of the templates and replacing the mutated AAs in the sequence. MODELLER is then run again to produce the mutant model.

#### Strengths

Our pipeline exploits a series of tools tailored to large sets of proteins. Useful information is provided at each step of the run so that decisions taken by the pipeline can be audited. In the case of a failure of the pipeline to provide a satisfactory structural model, the file *report.csv* traces the mutations on all the isoforms and provides an easy way to identify the isoform mapping the largest number of mutations. The previously mentioned *report.log* file is also important. This file contains all the data that would otherwise have to be manually collected such as the number of isoforms for a gene, the location of the mutations, which mutations can not be mapped on any known isoform and finally the values of the selection functions. These data can provide a useful start point if the user needs to manually model the protein. The user, upon data retrieval, can also decide if another isoform should be prioritised because of a mutation of particular interest not present in the isoform selected by the program, for example. Regarding the modelling part of the protocol, the final alignments, the used templates with detailed information on the selection process as well as the coverage are made available to the user. Although the process of building the variants can be time consuming if many of them need to be built, this part is fully automated. In the majority of the tested cases the models built are good and can be used for further studies (see Section Results). Thus, our pipeline reduces the time necessary to model a large number of proteins by automating the slowest parts of the process including the search for isoforms, the mapping of mutations, the search for crystallographic data to use as templates and the building of the alignments.

#### Limitations

As with any modelling study, also our method presents limitations. MoNvIso does not model the parts of the protein that are not covered by templates. The solution implemented in the program is the modelling of the single domains, although this implies the uncertainty on reciprocal orientations of the domains. An additional drawback is the possible presence of several small portions that can be modelled but are interspersed by regions not covered by templates. In some cases, the search for templates with HMMER does not return any result (depends on HMMER’s servers). When a lot of successive searches for homologues are queued on BLAST, a slowdown of the runs may occur. The coexistence of multiple mutations on the same protein is not taken into account by MoNvIso. The current version considers only single point mutations when building the variants.

#### Case study

We tested MoNvIso on a set of 68 proteins, evaluating 250 human isoforms extracted from the Uniprot database and relative mutations extracted from the relative Uniprot webpage, we capped the maximum number of mutations at 5. For all selected proteins MoNvIso was able to produce the alignments and to map the mutations onto the correct isoform. It successfully located and retrieved and edited the templates to generate the wild type (WT) models as well as, when the identified mutations are in the modelled portions, the variants.

Out of the 68 proteins we modelled, 51 WT models were compared to the ones available in the AlphaFold (AF) database (DB) (https://alphafold.ebi.ac.uk/), by extracting from the AF model the part of the sequence that we modelled and performing an RMSD analysis on the C*α*. For the remaining 17 proteins (BCL11A, CACNA1B, CAMKK1, CAMKK2, DNMT1, FMR1, GABRB3, GRIK2, GRM5, PLXNB1, SCN2A, SLC17A8, SNAP25, STX1A, SYN1, SYT1, TAF1), such comparison was not feasible because the isoform selected by MoNvIso was not the canonical one as considered by AF and the differences between the two isoforms lay in the regions we modelled. Thus, the number of C*α*s is different. It should be stressed that additionally for 13 proteins out of 68 we modelled an isoform different from the canonical sequence (thus, a total of 30 proteins out of 68 is associated with non-canonical isoforms) but the RMSD comparison with the AF models was possible because the changes were localised in region not covered by templates. The results of the comparison are presented in Table S1 together with the amount of residue for which AF has a high or very high confidence (pLDDT score > 70) about their position. The genes are ordered from the one with lower RMSD value to the highest. According to Table S1, 42 out of 55 (81%) models present an RMSD below 20 Å, and a visual inspection reinforces the validity of our results, since the larger RMSD values in this group are mainly due to small, disordered loops. The group of models with RMSD above 20 there are subunits assuming different orientation in our and AF structures. When comparing the number of AA with a high or very high confidence score, we see that in most of our results (44 out of 52), the portion that gets modelled retains at least 50% of these residues.

Two example structures are shown in Figure 2, for the proteins GRIN1 (Glutamate receptor ionotropic, NMDA 1; also known as GluN1; Uniprot #Q05586) and GRIN2B (Glutamate receptor ionotropic, NMDA 1; also known as GluN2B; Uniprot #Q13224). These two transmembrane proteins are subunits of the N-methyl-D-aspartate (NMDA) receptors glutamate receptors, which contribute to excitatory transmission in the brain. In the first case both AF and MoNvIso produce similar results that differ only in the domains for which no templates are available, but still modelled by AF. Examples are the C-terminal part, starting from K866 to S938 and the N-terminal helix (residues M1 to D23) that are modelled by AF and not by MoNvIso (see top left and bottom right in Figure 2a, respectively). These two portions of the models are removed from the model as a consequence of the MoNvIso protocol (see Step 3) since there are no templates to correctly model them, AF on the other hand always tries to model the whole chain. This leads to portions of the model with low or very low confidence scores (calculated by AF) and which corresponds to a pLDDT between 0 and 70, meaning that those parts of the model are unreliable.

**Figure 2.**
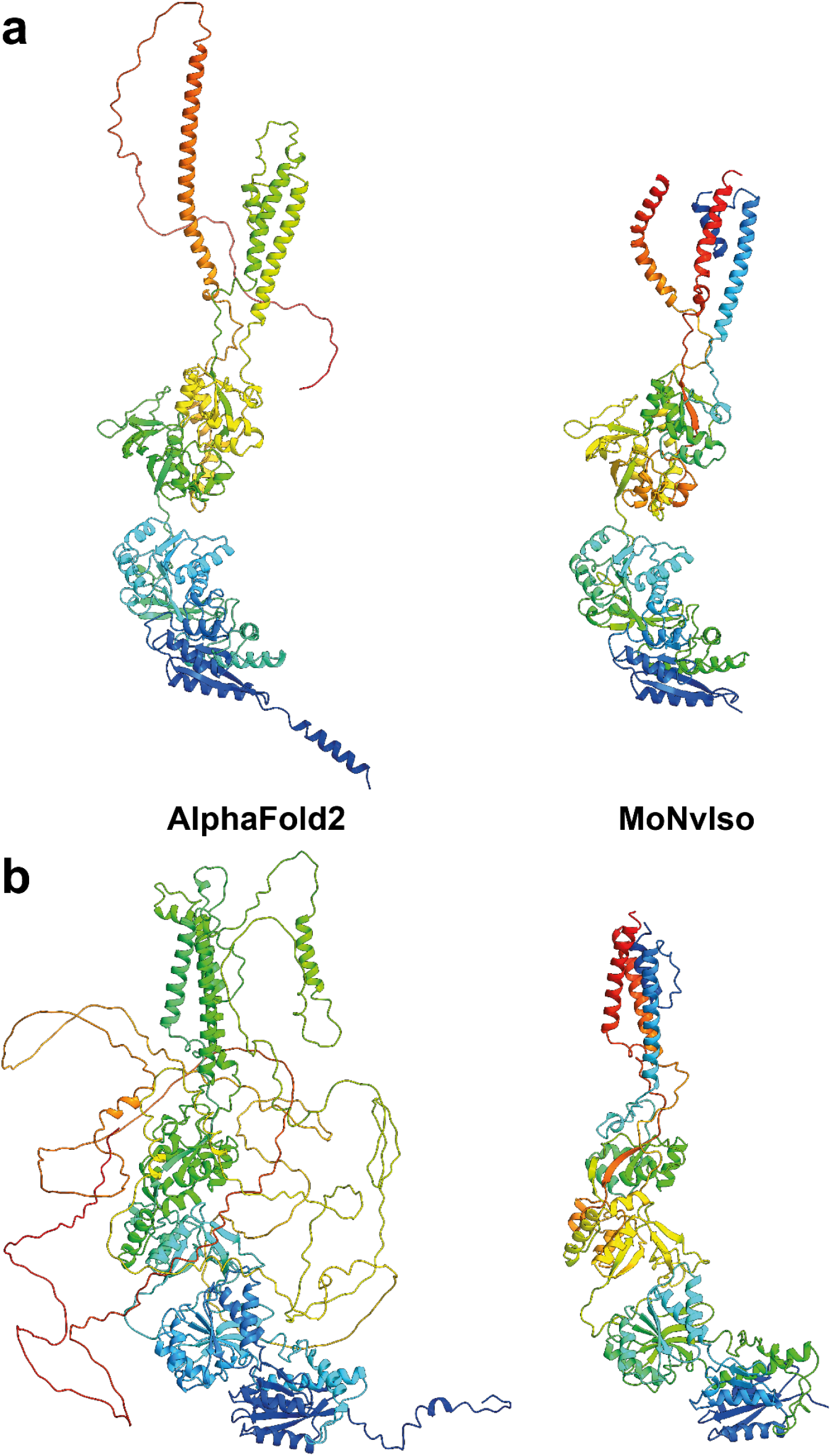
Comparison between the ribbon representations of GRIN1 (a) and GRIN2B (b) model structures generated with AF (left panels) and MoNvIso (right panels). The ribbons are colored from blue to red going from the N- to the C-terminal.

The results for GRIN2B demonstrate the differences between AF and MoNvIso predictions. AF successfully models the N-terminal part of the protein but fails to correctly build the trans and intra-membrane domains, which are then added as loops twisted around the correctly modelled section of the protein. Once again, the portions that are missing from the PDB database are poorly modelled. Since AF has been trained on the PDB dataset^9^ it still relies on available crystallographic data to correctly model structures. Thus, transmembrane domains such as those of GRIN2B, which are underrepresented in that training set because of the scarcity of experimentally determined structures of transmembrane proteins and their complexes^18^, may fail to be correctly built. On the other hand, MoNvIso automatically recognises which parts of the protein are possible to model with confidence and which are not and cuts the extra AAs out of the sequence, producing a model ready to be used for further analysis.

## Conclusions

Dissecting the impact of point mutations in the function of a protein are often hindered by a lack of an appropriate mapping of the mutation onto the correct isoform of a protein and the associated building of a reliable structure. Moreover, different isoforms of proteins can have widely differing functional roles and spatio-temporal expression profiles. As genomic variants associated with human traits and/or disease are being discovered at an increasing rate, approaches to modelling them are urgently needed. Our computational protocol MoNvIso addresses these two aspects; being able to select the correct isoform of a protein containing the mutations of interest and to largely automate the construction of the associated protein structure. Although platforms that provide accurate structures of a protein are available and routinely used, surprisingly few of them can be implemented in a pipeline to automate the modelling of multiple different proteins. Therefore, our protocol combines this final step with the key preliminary assessment of the isoform mapping correctly the mutation of interest.

The test of MoNvIso on a set of proteins and the comparison with the results of AlphaFold confirms the validity of our approach. Additionally, our computational protocol can be easily inserted in a pipeline suitable to perform extensive campaigns of investigation on protein-protein interactions. MoNvIso is particularly useful to evaluate the availability of templates for large sets of proteins and automatically selecting the correct isoform based on the mutations of interest. MoNvIso is freely available and can be downloaded from github at the following link: https://github.com/MoNvIsoModeling/MoNvIso, implemented in Python from 3.8 and tested on version 2.7, 3.0, 3.7 and 3.9 and supported on Linux.

## Acknowledgements

PC acknowledges the Deutsche Forschungsgemeinschaft (DFG) via the Research Training Group RTG2416 MultiSenses-MultiScales (368482240/GRK2416).

Silvia De Rubeis received a Wilhelm Bessel Research Award from the Alexander von Humboldt Foundation.

We thank Dr. Emiliano Ippoliti (Jülich) and Dr. Enrico Gandini (Milan) for technical support. JDA and OS received funding from the European Union’s Horizon 2020 Framework Programme for Research and Innovation under the Specific Grant Agreement Nos. 945539 (Human Brain Project SGA3).

## Required Python 3.0 libraries

-**Biopython** to edit PDB files (https://biopython.org/wiki/Download)

