## Supplementary Materials for "Modelling eNvironment for Isoforms (MoNvIso): A general platform to predict structural determinants of protein isoforms in genetic diseases"

In Table S1 we compare the MoNvIso structural models with the structures extracted from the AlphaFold2 (AF) database[1].

**Table S1:**
Comparison between MoNvIso results and AF The pLDDT is a measurement, used by AF, of the confidence of the position of each residue in the predicted model. The gene labels are taken from the Uniprot[2] database.

| GENE | RMSD (Å) | AF C$\alpha$ with pLDDT > 70* | MoNvIso C$\alpha$ with pLDDT > 70 | MoNvIso residues (pLDDT > 70) / AF residues (pLDDT > 70) (%) | MoNvIso Modeled residues | AF Modeled residues |
|---|---|---|---|---|---|---|
| GAD1 | 0.6 | 503 | 502 | 99.80 | 502 | 594 |
| PVALB | 0.8 | 110 | 108 | 98.18 | 108 | 110 |

| | | | | | | |
|---|---|---|---|---|---|---|
| PTCH1 | 0.9 | 1007 | 1003 | 99.60 | 1028 | 1447 |
| EFNB1 | 1.2 | 186 | 141 | 75.81 | 141 | 346 |
| NLGN4X | 1.3 | 586 | 539 | 91.98 | 565 | 816 |
| KDM6A | 1.4 | 871 | 474 | 54.42 | 487 | 1401 |
| BDNF | 1.7 | 140 | 106 | 75.71 | 114 | 247 |
| PTPN11 | 1.8 | 504 | 504 | 100.00 | 531 | 593 |
| RIMS1 | 2.0 | 486 | 140 | 28.81 | 150 | 1692 |
| EIF4E | 2.1 | 188 | 188 | 100.00 | 191 | 217 |
| RAB3A | 2.4 | 176 | 176 | 100.00 | 178 | 220 |
| GPHN | 2.6 | 592 | 419 | 70.78 | 419 | 736 |
| KIF11 | 2.7 | 757 | 314 | 41.48 | 355 | 1056 |
| CACNG2 | 2.9 | 150 | 150 | 100.00 | 216 | 323 |
| KIF5C | 2.9 | 776 | 332 | 42.78 | 350 | 957 |
| NSD1 | 2.9 | 690 | 223 | 32.32 | 237 | 2696 |
| GRIN2B | 3.0 | 694 | 694 | 100.00 | 801 | 1484 |
| CDKL5 | 3.4 | 277 | 277 | 100.00 | 307 | 960 |
| CTNND2 | 3.5 | 456 | 413 | 90.57 | 465 | 1225 |
| RAC1 | 3.6 | 179 | 179 | 100.00 | 189 | 192 |
| MECP2 | 3.9 | 77 | 69 | 89.61 | 77 | 486 |
| NTF4 | 4.1 | 131 | 112 | 85.50 | 121 | 210 |
| GRIN1 | 4.2 | 778 | 761 | 97.81 | 808 | 938 |
| STXBP1 | 4.2 | 543 | 543 | 100.00 | 594 | 594 |
| CDH2 | 4.4 | 668 | 543 | 81.29 | 559 | 906 |
| AP2M1 | 4.6 | 409 | 409 | 100.00 | 435 | 435 |
| APP | 5.6 | 427 | 156 | 36.53 | 176 | 770 |
| FOXP1 | 5.7 | 196 | 81 | 41.33 | 89 | 677 |
| DDX3X | 5.9 | 407 | 404 | 99.26 | 448 | 662 |
| GRM2 | 6.3 | 765 | 759 | 99.22 | 791 | 872 |
| OPA1 | 6.7 | 692 | 306 | 44.22 | 331 | 960 |
| HOMER1 | 7.5 | 292 | 117 | 40.07 | 143 | 354 |
| PTEN | 8.0 | 315 | 311 | 98.73 | 349 | 403 |
| STX1B | 8.8 | 237 | 209 | 88.19 | 257 | 288 |
| CREBBP | 10.7 | 823 | 581 | 70.60 | 645 | 2442 |

| | | | | | | |
|---|---|---|---|---|---|---|
| SLC1A3 | 10.8 | 429 | 414 | 96.50 | 458 | 542 |
| EP300 | 11.6 | 833 | 582 | 69.87 | 649 | 2414 |
| TBR1 | 13.4 | 176 | 176 | 100.00 | 234 | 682 |
| APBA2 | 14.0 | 322 | 229 | 71.12 | 294 | 749 |
| UBE3A | 15.9 | 702 | 413 | 58.83 | 429 | 875 |
| DRD2 | 16.4 | 271 | 271 | 100.00 | 377 | 443 |
| GRIA1 | 17.0 | 738 | 734 | 99.46 | 787 | 906 |
| DYRK1A | 21.2 | 373 | 373 | 100.00 | 426 | 763 |
| GRIA2 | 23.2 | 763 | 755 | 98.95 | 787 | 883 |
| ATRX | 23.4 | 843 | 542 | 64.29 | 778 | 2492 |
| EPHB2 | 26.4 | 835 | 765 | 91.62 | 834 | 1055 |
| CTCF | 29.9 | 304 | 301 | 99.01 | 323 | 727 |
| FRMPD4 | 31.8 | 441 | 368 | 83.45 | 441 | 1322 |
| GAP43 | 33.7 | 31 | 31 | 100.00 | 170 | 238 |
| DLGAP2 | 42.6 | 148 | 107 | 72.30 | 344 | 1054 |
| UNC13A | 52.6 | 1164 | 974 | 83.68 | 1099 | 1703 |
| PTPRT | 69.4 | 1241 | 1111 | 89.52 | 1153 | 1441 |

**Example**
For the sake of clarity on the use of MoNvIso, we outline a scheme of the input and output files of a typical run, including a more detailed description of the workflow of the code.

As input, a list of genes and mutations is provided. We consider GRIN1 and GRIN2B a along with 6 mutations per gene. The mutations are inserted in the possible notations accepted by the code, i.e., single letter, three letters, separated strings, single string:

```
GRIN1
R	844	C
Ala	349	Thr
Ala	645	Ser
Pro	578	Arg
Ser	688	Tyr
Tyr	647	Ser

GRIN2B
E413G
C436R
M1342R
```

S1415L
L1424F
P1439A

Note that a blank line separates the groups of data associated with each gene and that no blank line is to be present at the end of the input file.

For each gene and isoform in the input file a folder is created where all the files will be stored.

According to Uniprot, there are 8 isoforms of GRIN1 and only one of GRIN2B. All the isoforms of each gene are evaluated one by one. In this example we will focus our attention on the isoform 1 of GRIN1.

First, the code searches for homologous sequences through the NCBI BLAST[3] application programming interface (API), which are saved in a file labelled with the name of the isoform and "_hits.fasta", "isoform1_hits.fasta" in our case. The sequences are then aligned using COBALT[4] and the multi-sequence alignment (MSA) is stored in the file "homologous_aligned.fasta". The alignment file is used to build the Hidden Markov Model (HMM) by HMMER version 3.3.2 (http://hmmer.org/), the file containing the HMM is named according to the gene name ".hmm", "GRIN1.hmm" in our case. These files can be found in the "template_search" folder.

Successively, the HMM is used to identify templates using the HMMER API, the result of the search is stored in a file called "possible_templates.xml", and, if available, the most similar *N* templates are selected, where *N* is specified by the user using the "parameters.dat" file with the keyword "PDB_TO_USE". If the resolution of the templates is below the threshold selected by the user (using the RESOLUTION keyword in the parameter file), the desired chain(s) are extracted and aligned to the MSA, again with COBALT. A file containing only standard amino acids (AAs) without disordered atoms is produced and called as the PDB "identifier_chain_cleaned.pdb", for example "6whr_A_clean.pdb" in the case of GRIN1. The sequence identity, the coverage and the intervals of the target sequence covered by each template are calculated and reported in the "covered_intervals" file. The "Score" reported in the covered_intervals file is the number of AAs of the target sequence aligned with AAs of the template, while the "Sequence Identity" is the portions of aligned AAs that are also identical. These files can be found in the "template_analysis" folder, the pdb files will be stored in the "pdbs" folder.

In the gene folders other files are also generated:

- The "master_isoforms.txt" file contains the name of all the possible isoform files.
- The "usable_isoforms.txt" files contains the isoforms that could be evaluated and did not occur in errors or absence of valid templates.
- The files called "isoform0.fasta" always contains the sequence of the canonical isoform.

Each isoform folder will contain:

- a file called “final_msa.fasta” containing the sequences of the downloaded PDBs aligned to the MSA.
- The file “alignment_pdbs.fasta” is a subset of the “final_msa.fasta” containing only the PDBs and the target sequence.
- The “aligned_oneline_withgaps.fasta” is the same as “alignment_pdbs.fasta” but the sequence is written as one line, which is easier to read for the algorithm.
- The PDB files that were selected from the filtering process (see Manuscript).

Once the appropriate isoform is selected,  the model of the wild type is generated by MODELLER[5] and stored in the folder “modeller_files_wt”. In the file MYOUT.dat the model with the best DOPE[6] score is indicated. This model will be kept in the folder “modeller_files_wt” while other models possibly generated by MODELLER but with lower DOPE scores will be moved in the “models” subfolder.
The same procedure will be carried out for the variants, being the results collected in folders called mutation_MUTATIONLABEL, “mutation_A349T” in our case by considering the A349T mutation of GRIN1.
The number of models produced by MODELLER can be specified using the keyword “NUM_OF_MOD_WT”, and the number of model of mutant using the keyword “NUM_OF_MOD_MUT”, the user can also specify the number of not-covered AA necessary to cut the protein with the “MODEL_CUTOFF” keyword.

The folder containing the files of the modelled isoform also contains the file necessary to run modeller, namely “run_modeller.py” and “input_modeller.dat” that contain the python code and the alignments, respectively. The file “for_modeller” contains the template sequence aligned to the MSA. The file “modeller_output.txt” contains all the messages that modeller would normally print on screen, so that it is possible to keep track if something goes wrong.

For each gene two reports are generated: one called report.csv shows where the mutations are located, the other (report.log) collects the isoform selected by MoNvIso, how many and which mutations are mapped on each isoform, how many and which mutations are in modellable regions, which mutations that are not mapped on any isoform, and the scores both partials and total achieved by each isoform (see Equations 1-3 in the Manuscript). If more than one gene is considered, a final report is also produced by pasting all single report.log.


1. Tunyasuvunakool, K. *et al.* Highly accurate protein structure prediction for the human proteome. *Nature* **596**, 590–596 (2021).
2. Bateman, A. *et al.* UniProt: the universal protein knowledgebase in 2021. *Nucleic Acids Res.* **49**, D480–D489 (2020).
3. Basic local alignment search tool - ScienceDirect. https://www.sciencedirect.com/science/article/pii/S0022283605803602.
4. COBALT: constraint-based alignment tool for multiple protein sequences | Bioinformatics |

Oxford Academic.
https://academic.oup.com/bioinformatics/article/23/9/1073/272774?login=true.

5. Webb, B. & Sali, A. Comparative Protein Structure Modeling Using MODELLER. *Curr. Protoc. Bioinforma.* **54**, (2016).
6. Shen, M. & Sali, A. Statistical potential for assessment and prediction of protein structures. *Protein Sci.* **15**, 2507–2524 (2006).
